## Supplementary material for "N-terminal VP1 truncations favor *T*=1 norovirus-like particles": complete supplement


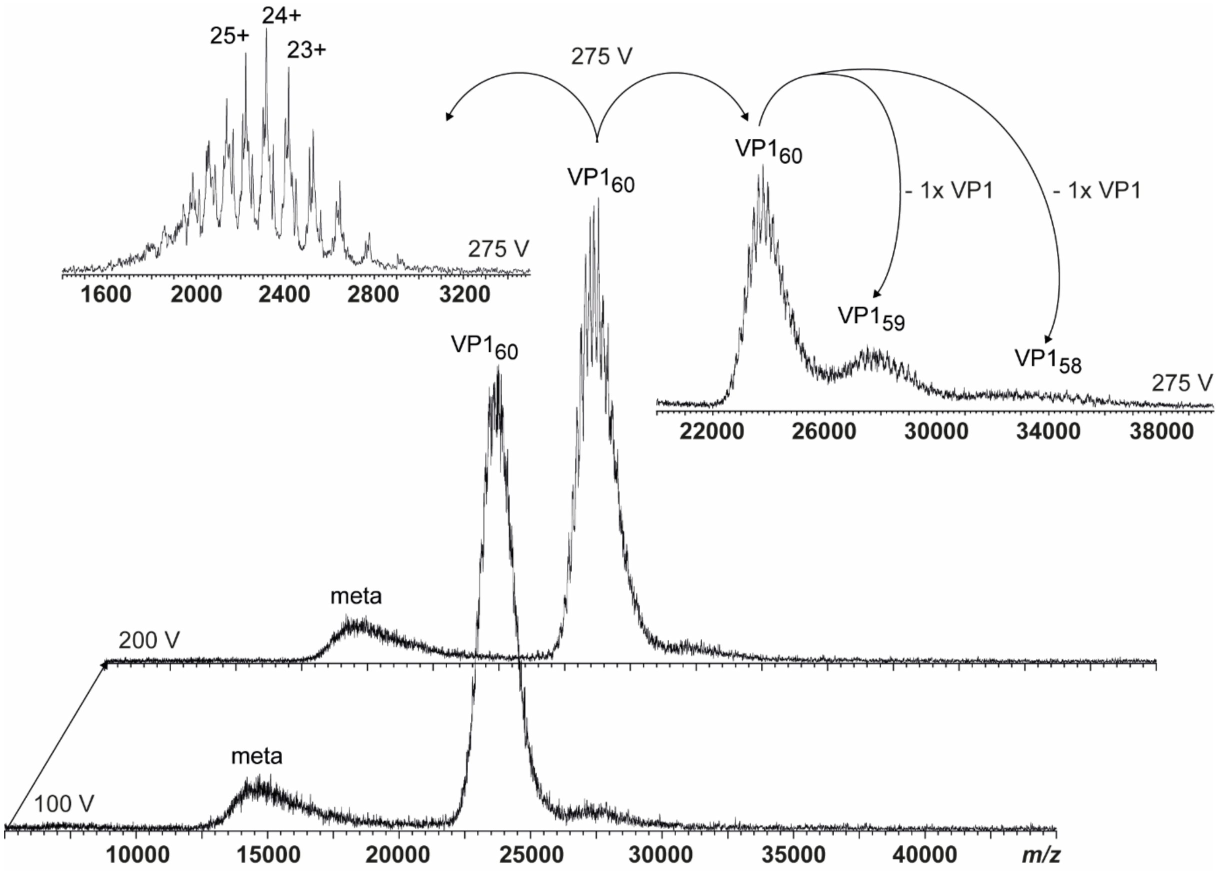


**Figure S1.** Native MS of a GII.10 Vietnam VLPs. Dissociation pathway without selection in the quadrupole shown for GII.10 Vietnam in 250 mM ammonium acetate pH 7 at 10 µM VP1. From bottom to top, illustrative mass spectra are shown for 100 V, 200 V, and a zoom in to low *m*/*z* (monomer) and high *m*/*z* (VP1 60-, 59-, 58-mer) at 275 V acceleration into the collision cell. As lower mass ions at approximately 15,000 *m*/*z* are annotated as metastable ions (meta), monomer lacking at least 45 aa most likely dissociate from *T*=1 species.

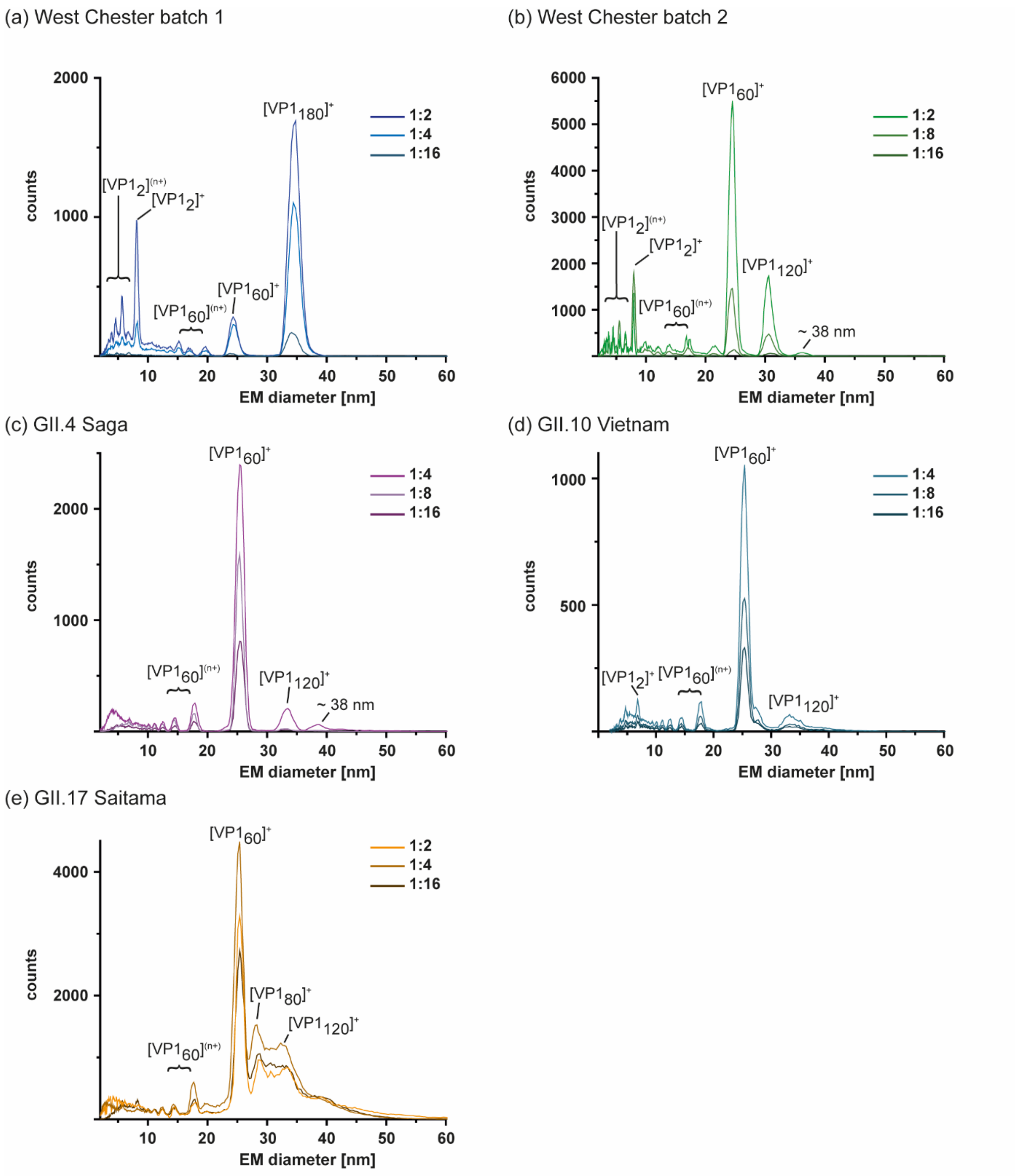


**Figure S2.** GEMMA spectra of different hNoVLPs in 40 mM ammonium acetate at pH 7, 3 different tested dilutions exemplarily shown for (a) West Chester batch 1 (b) West Chester batch 2 (c) GII.4 Saga (d) GII.10 Vietnam and (e) GII.17 Saitama. In GI.1 West Chester batch 2 and GII.4 Saga, additional species at approximately 38 nm were assigned to aggregation due to their appearance only at higher concentrations.

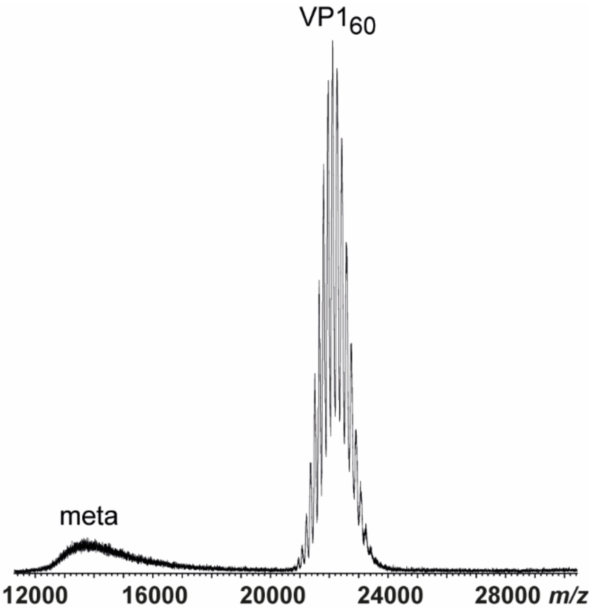


**Figure S3.** Native mass spectrum of GII.4 Saga VLPs at 50 mM ammonium pH 9 at 10 μM VP1. In contrast to GEMMA measurements (Figure 5), T=1 particles are detected.

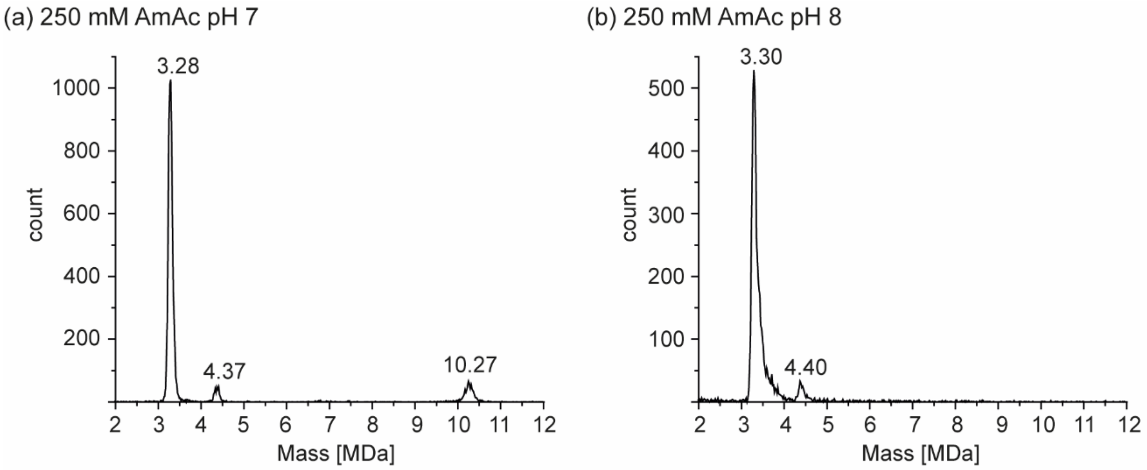


**Figure S4.** Charge detection mass spectra of GI.1 West Chester batch 1 VLPs in 250 mM ammonium acetate at (a) pH 7 and (b) pH 8. Next to *T*=1 particles, a species at approximately 4.37 MDa at pH 7 and 4.40 MDa at pH 8 is detected. Given the VP1 60-er mass 4.4 MDa species equal VP1 80-mers. At pH 7, further ions at 10.27 MDa or *T*=3 particles are observed.

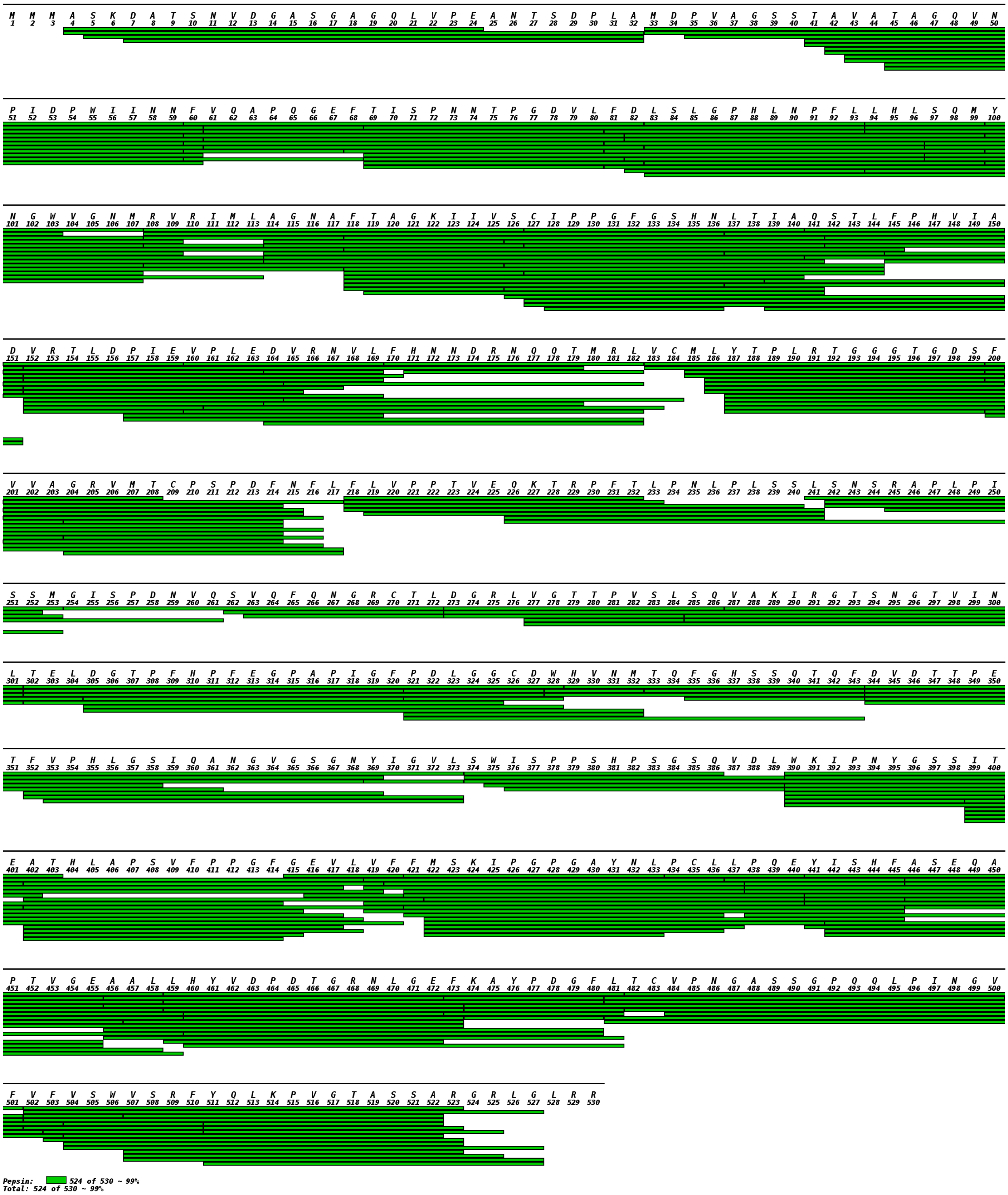


**Figure S5.** VP1 mapping overview of hNoVLP GI.1 West Chester batch 1 after pepsin digestion. In total 524 of 530 aa are covered (coverage 99 %). The first N-terminal residue covered is Ala4, while C-terminally the triplet Leu528-Arg529-Arg530 is missing.

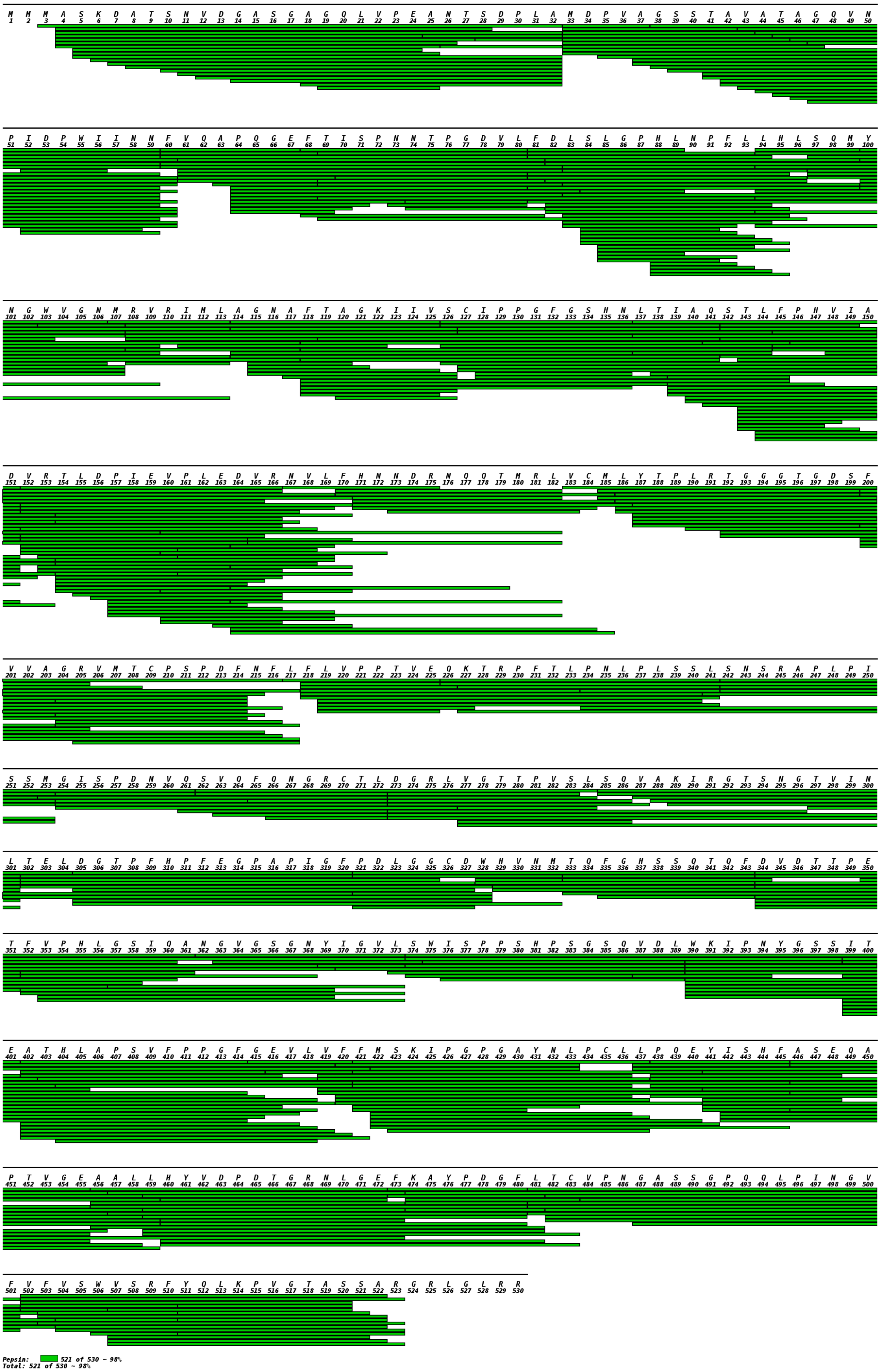


**Figure S6.** VP1 mapping overview of hNoVLP GI.1 West Chester batch 2 after pepsin digestion. In total 521 of 530 aa are covered (coverage 98 %). The first N-terminal residue covered is Met3 and the C-terminus is covered up to residue Arg523.

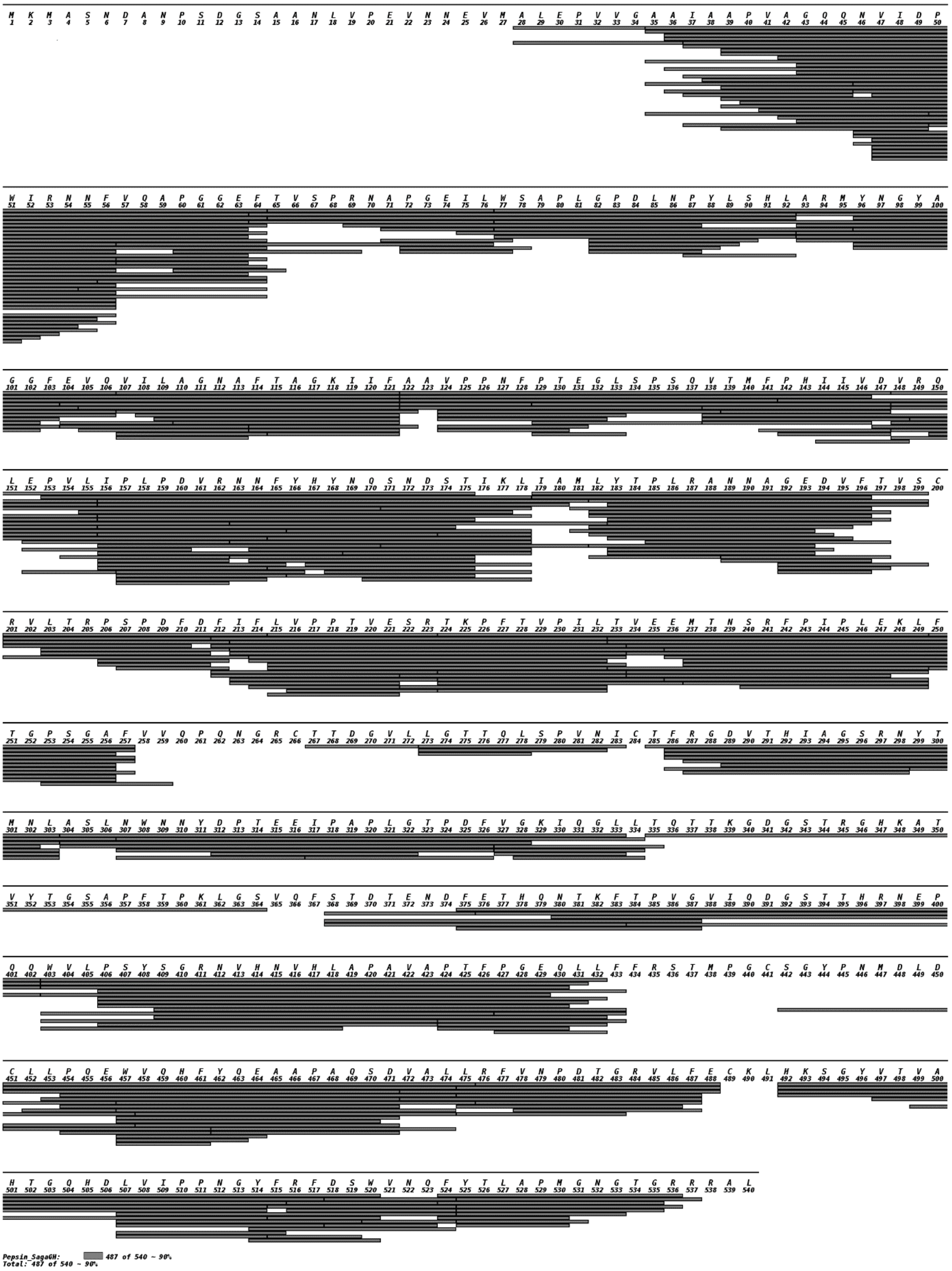


**Figure S7.** VP1 mapping overview of hNoVLP GII.4 Saga after pepsin digestion. In total 487 of 540 aa are covered (coverage 90 %). The first N-terminal residue covered is Ala28, while C-terminally the triplet Arg538-Ala539-Leu540 is not covered.

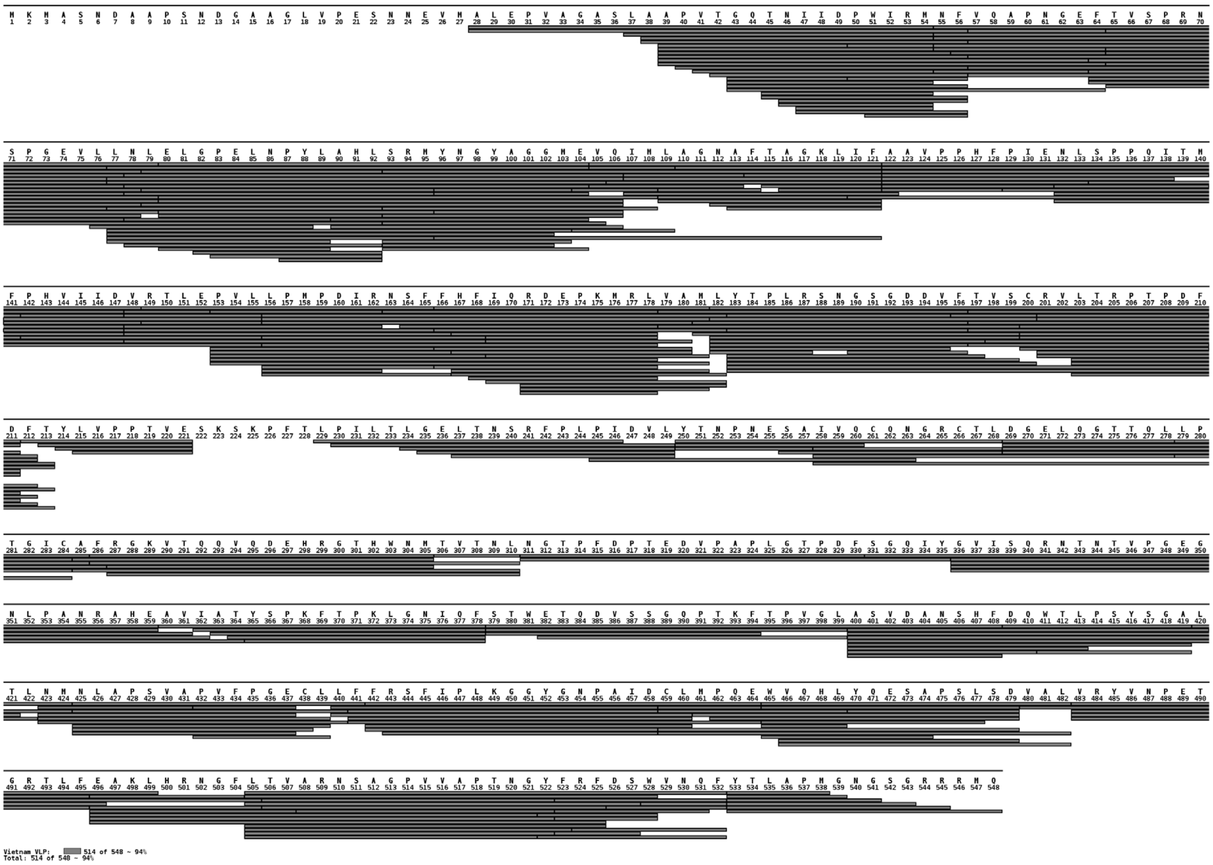

**Figure S8.** VP1 mapping overview of hNoVLP GII.10 Vietnam after pepsin digestion. In total 514 of 548 aa are covered (coverage 94 %). The first N-terminal residue covered is Ala28, while full-coverage is given for the C-terminus.

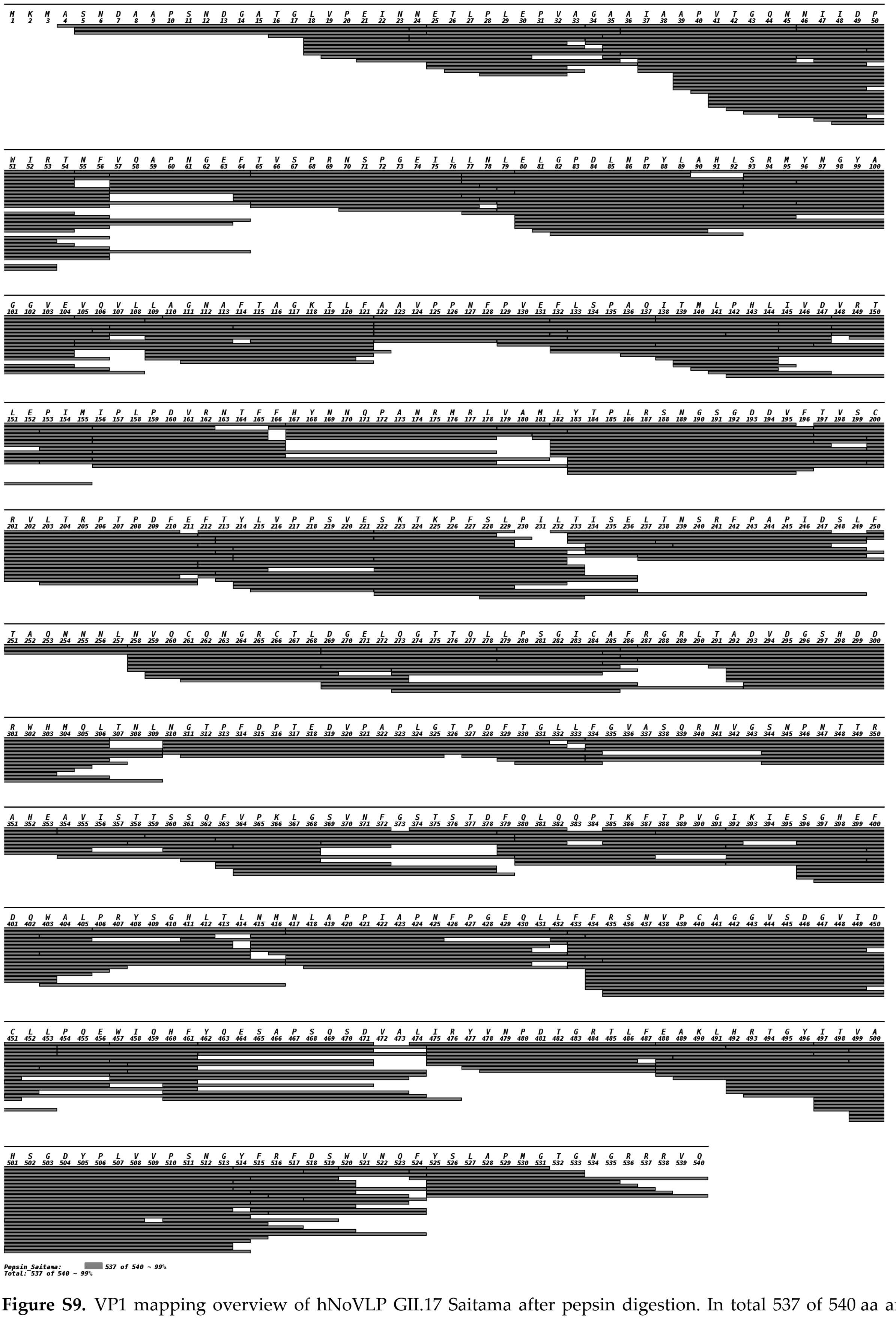

**Figure S9.** VP1 mapping overview of hNoVLP GII.17 Saitama after pepsin digestion. In total 537 of 540 aa are covered (coverage 99 %). The first N-terminal residue covered is Ala4, while full-coverage is given for the C-terminus.

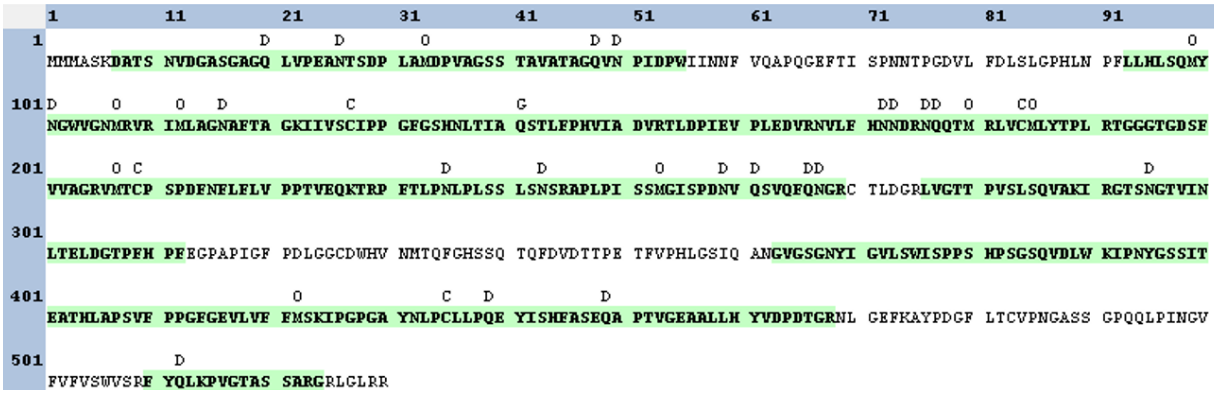


**Figure S10.** VP1 mapping overview of hNoVLP GI.1 West Chester batch 1 after trypsin digestion (coverage 72 %). The first N-terminal residue covered is Asp7 and C-terminally six residues are not covered.

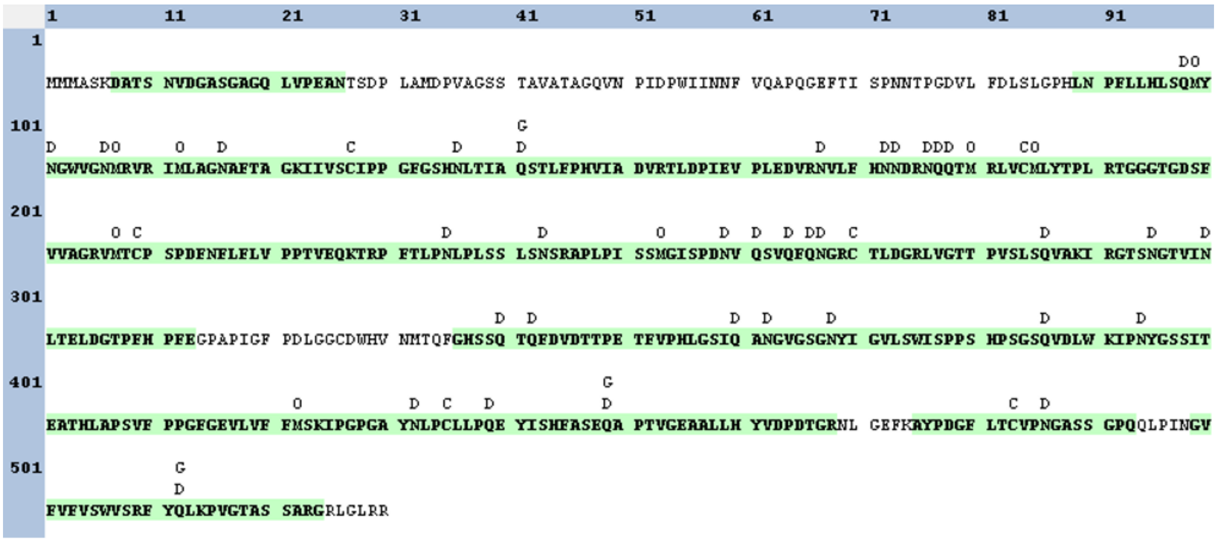


**Figure S11.** VP1 mapping overview of hNoVLP GI.1 West Chester batch 2 after trypsin digestion (coverage 80 %). The first N-terminal residue covered is Asp7 and C-terminally six residues are not covered.

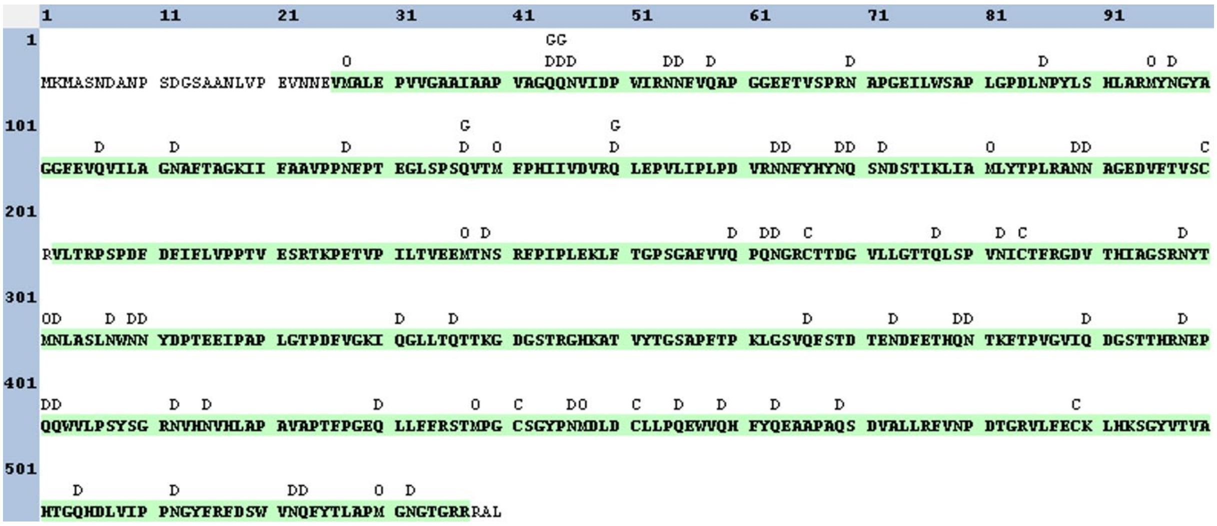


**Figure S12.** VP1 mapping overview of hNoVLP GII.4 Saga after trypsin digestion (coverage 95 %). Semi-tryptic peptide search identifies Val26 as the first N-terminal residue covered, while C-terminally the triplet Arg538-Ala539-Leu540 is not covered.

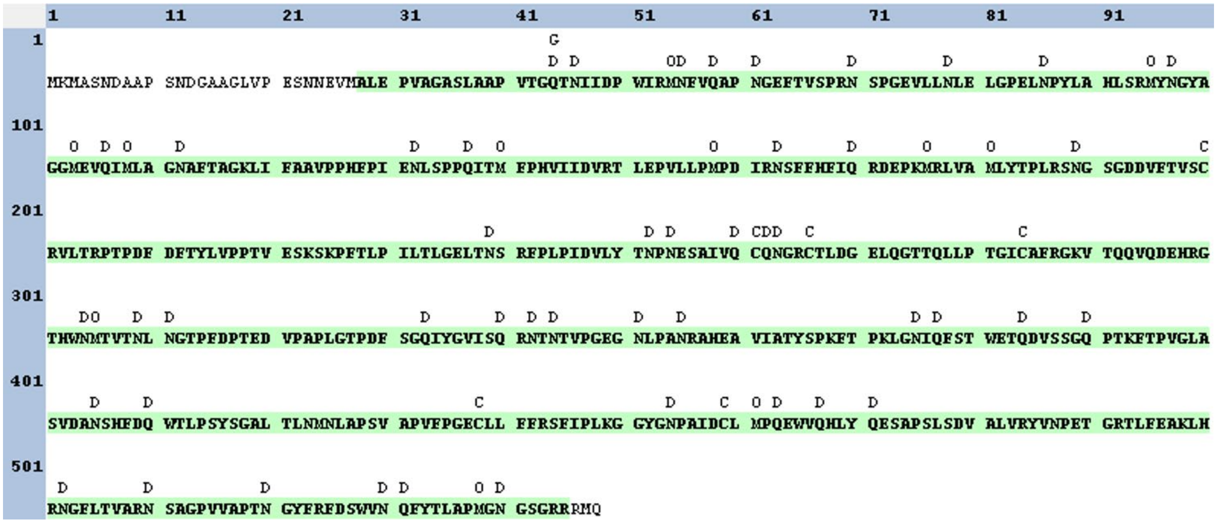

**Figure S13.** VP1 mapping overview of hNoVLP GII.10 Vietnam after trypsin digestion (coverage 95 %). Semi-tryptic peptide search identifies Ala28 as the first N-terminal residue covered, while C-terminally the triplet Arg546-Met547-Gln548 is not covered. Note that the sequence covered starts indeed with a semi-tryptic peptide.

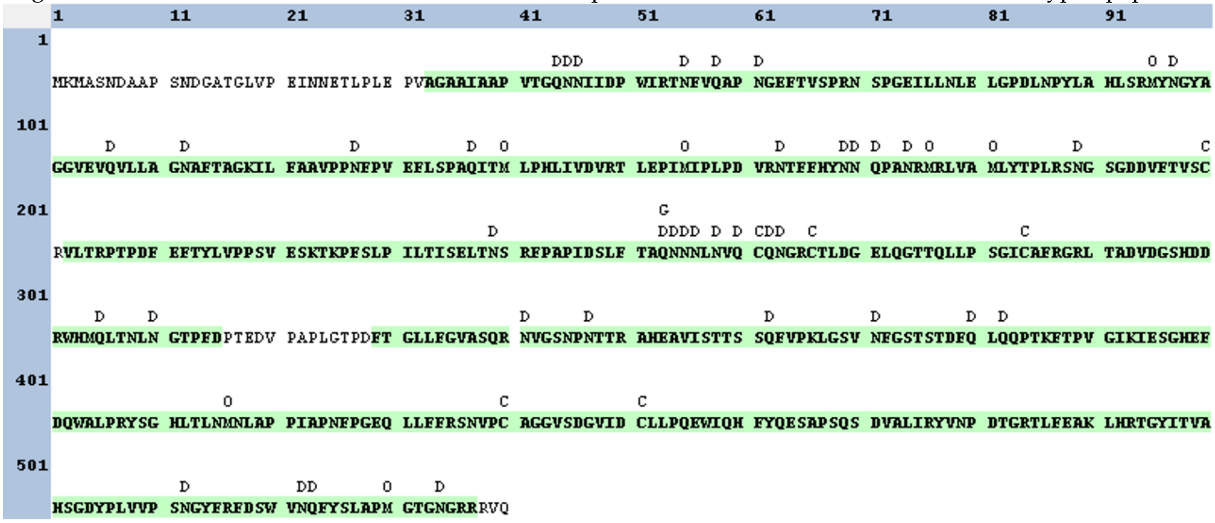

**Figure S14.** VP1 mapping overview of hNoVLP GII.17 Saitama after trypsin digestion (coverage 91 %). Semi-tryptic peptide search identifies Ala33 as the first N-terminal residue covered, while C-terminally the triplet Arg538-Val539-Gln540 is not covered. Note that the sequence covered starts indeed with a semi-tryptic peptide.

**Table S1.** Data mass table for charge detection mass spectrometry measurements. Abbreviations Th. theoretical, Exp. experimental, Calc. calculated. VP1 Calc. based on VP1 60-mer mass. For VP1 Th. and VP1 Calc. the total aa-amount and aa truncation according to experimental mass are given, respectively. *N* VP1 given for larger assemblies respective to VP1 Calc. *indicate masses are approximations due to low particle counts.

| Variant | VP1 Th. | Complex Exp. | VP1 Calc. | *N* VP1 Complex |
|---|---|---|---|---|
| GII.4 Saga | 59005 Da 540 aa | 3.35 MDa | 55800 Da, -31 aa | 60 |
| GII.10 Vietnam | 59901 Da 548 aa | 3.41 MDa | 56800 Da, -31 aa | 60 |
| | | ~ 4.5 MDa* | | 79 |
| | | ~ 6.9 MDa* | | 121 |
| GII.17 Saitama | 58957 Da 540 aa | 3.44 MDa | 57300 Da, -17 aa | 60 |
| | | 4.07 MDa | | 71 |
| | | 5.20 MDa | | 91 |
| | | 5.72 MDa | | 100 |
| | | 6.20 MDa | | 108 |
| | | 6.87 MDa | | 120 |

**Table S2.** Data mass table for conventional QToF measurements. Abbreviations Th. theoretical, Exp. experimental, Calc. calculated. For all tested variants, collision induced dissociation experiments resulted in dissociated VP1 monomer with a main species (VP1 Exp. main) and one or two neighboring species with lower intensity (VP1 Exp. following). VP1 60-mer experimental mass (Complex $VP1_{60}$ Exp.) assigned where charge state resolution was obtained and given as approximations for GI.1 West Chester due to low desolvation and associated low charge state resolution. *approximation.

| Variant | VP1 Th. | VP1 Exp. main | VP1 Exp. following | Complex $VP1_{60}$ Exp. |
|---|---|---|---|---|
| GI.1 West Chester | 56609 Da, 530 aa | 52760 ± 10 Da, -40 aa | 52540 ± 10 Da, -43 aa- | ~3.4 MDa* |
| GII.4 Saga | 59005 Da 540 aa | 54600 ± 20 Da, -45 aa | 54270 ± 20 Da, -48 aa | 3.27 ± 0.02 MDa |
| GII.10 Vietnam | 59901 Da 548 aa | 55560 ± 10, -45 aa | 55220 ± 20, -48 aa<br>56290 ± 10, -37aa | 3.33 ± 0.02 MDa |

**Table S3 GEMMA** Data mass table for gas phase electrophoretic molecular mobility analysis. Abbreviations, Exp. experimental, Calc. calculated. VP1 oligomers given as experimental EMD values and respective molecular weight calculations for low EMD range ([1] Bacher et al 2001) and high EMD range ([2] Weiss et al. 2019) *approximation due to low particle counts.

| Variant | pH | VP1 dimer Exp. (nm) Calc.[1] (kDa) | VP1 60-mer Exp. (nm) Calc.[2] (MDa) | VP1 80-mer Exp. (nm) Calc.[2] (MDa) | VP1 120-mer Exp. (nm) Calc.[2] (MDa) | VP1 180-mer Exp. (nm) Calc.[2] (MDa) |
|---|---|---|---|---|---|---|
| GI.1 West Chester batch 1 | 6 | | 24.29 ± 0.09<br>3.37 ± 0.03 | | | 34.54 ± 0.05<br>8.50 ± 0.03 |
| | 7 | 8.10 ± 0.05<br>112 +2 | 24.09 ± 0.27<br>3.23 ± 0.10 | | | 34.37 ± 0.13<br>8.40 ±0.09 |
| | 9 | 8.03 ± 0.01<br>109 ±1 | | | | |
| GI.1 West Chester batch 2 | 6 | | 24.48 ± 0.02<br>3.44±0.01 | 30.73 ± 0.05<br>6.25±0.03 | | |
| | 7 | 8.00 ± 0.07<br>108 ± 3 | 24.50 ± 0.12<br>3.45±0.04 | 30.71 ± 0.17<br>6.24±0.09 | | |
| | 9 | 7.89 ± 0.01<br>104 ± 1 | 24.18 ± 0.06<br>3.33 ± 0.02 | | | |
| GII.4 Saga | 5 | | 25.27 ± 0.01<br>3.74 ± 0.01 | | | |
| | 6 | | 25.18 ± 0.01<br>3.70±0.01 | | | |
| | 7 | | 25.38 ± 0.07<br>3.8±0.03 | | 33.30 ± 0.08<br>7.72 ± 0.05 | |
| | 8 | 8.03 ± 0.02<br>109 ± 1 | 25.44 ±0.03<br>3.80 ± 0.01 | | | |
| | 9 | 7.88 ± 0.01<br>104 ± 1 | | | | |
| GII.10 Vietnam | 5 | | ~ 25* | | | |
| | 6 | | 25.11 ± 0.01<br>3.68±0.02 | | | |
| | 7 | | 25.32 ± 0.02<br>3.75 ± 0.01 | | 33.41 ± 0.09<br>7.79 ± 0.05 | |
| | 8 | 7.90 ± 0.02<br>104 ± 1 | 25.10 ± 0.02<br>3.67 ± 0.01 | | 33.31 ± 0.05<br>7.73 ± 0.03 | |
| | 9 | 7.93 ± 0.02<br>105 ± 1 | 25.06 ± 0.02<br>3.66 ± 0.01 | | | |
| GII.17 Saitama | 5 | | 25.47 ± 0.09<br>3.81 ± 0.04 | 28.67 ± 0.54<br>5.21 ±. 0.03 | | |
| | 6 | | 25.20 ± 0.06<br>3.71 ± 0.03 | | | |
| | 7 | | 25.36 ± 0.06<br>3.77 ± 0.03 | 28.48 ± 0.17<br>5.1 2± 0.08 | 32.62 ± 0.22<br>7.31 ± 0.13 | |
| | 8 | 7.99 ± 0.08<br>108 ± 3 | 25.14 ± 0.05<br>3.69 ± 0.02 | 28.13 ± 0.10<br>4.9 6± 0.05 | 32.33 ± 0.08<br>7.15 ± 0.05 | |
| | 9 | 8.08 ± 0.05<br>111 ± 2 | 25.22 ± 0.17<br>3.72 ± 0.07 | | | |

[1] Bacher G, Szymanski WW, Kaufman SL, Zollner P, Blaas D, Allmaier G. Charge-reduced nano electrospray ionization combined with differential mobility analysis of peptides, proteins, glycoproteins, noncovalent protein complexes and viruses. J Mass Spectrom. 2001;36(9):1038-52.

[2] Weiss VU, Pogan R, Zoratto S, Bond KM, Boulanger P, Jarrold MF, et al. Virus-like particle size and molecular weight/mass determination applying gas-phase electrophoresis (native nES GEMMA). Anal Bioanal Chem. 2019;411(23):5951-62
